# Integrated multiomics reveals inflammation-driven excessive erythrocytosis in subjects with Monge’s disease

**DOI:** 10.1101/2025.10.20.683487

**Authors:** Priti Azad, Andrew B. Caldwell, Francisco C. Villafuerte, Shamieh Banihani, Dan Zhou, Shankar Subramaniam, Gabriel G. Haddad

## Abstract

Monge’s disease, or Chronic Mountain Sickness (CMS), is a chronic high-altitude disorder characterized by hypoxia-induced excessive erythrocytosis (EE), elevating the risk of stroke and myocardial infarction. Using RNA-seq and ATAC-seq, we profiled iPSC-derived erythroid cells from CMS and non-CMS subjects under normoxia and hypoxia to identify statistically significant, disease-associated transcriptional and chromatin accessibility changes. RNA-seq revealed induction of inflammatory, stress, and erythropoiesis programs in CMS even under normoxia, including robust activation of JAK/STAT signaling, upregulation of heme metabolism and VEGF, and accelerated erythrocyte lineage commitment alongside repression of Notch and WNT/β-catenin. Hypoxia amplified this dysregulated state, and critically, activated NFκB-driven inflammatory signaling together with canonical HIF targets. ATAC-seq revealed pronounced hypoxia-induced changes, with increased accessibility within inflammatory and erythrocyte lineage genes occurring concomitantly with decreased accessibility within pluripotency and ectodermal lineage genes. Pharmacological NFκB inhibition in CMS cells significantly reduced EE (*p*-value <0.0001), whereas NFκB activation in non-CMS cells was sufficient to drive EE (*p*-value <0.01), confirming the causal role inferred by our multiomics analyses. Collectively, our multiomics and functional experiments substantiate a coordinated chromatin–transcription paradigm favoring an inflammatory axis that, through hypoxia-driven NFκB activation, accelerates stress-induced erythroid commitment and underlies EE in CMS.

## Introduction

Chronic Mountain Sickness (CMS), also known as Monge’s disease, is a pathological response to chronic high altitude hypoxia, particularly prevalent among populations like the long-term residents of the Andean city of Cerro de Pasco, Peru^1-4^ (>4300m). Its defining feature is excessive erythrocytosis (EE), marked by hemoglobin levels ≥21g/dL in men and ≥19g/dL in women. This overproduction of red blood cells elevates blood viscosity, reduces tissue perfusion, and increases the risk of stroke and myocardial infarction leading to early mortality, especially in males^3^. Notably, a subset of individuals living at the same altitude remain unaffected: these non-CMS individuals can endure high altitude hypoxia while maintaining normal erythropoietic function. Comparative investigation of genetic and epigenetic mechanisms governing lineage commitment, erythroid differentiation, proliferation, and apoptosis under chronic hypoxia in CMS and non-CMS cells offers a unique opportunity to uncover critical mechanisms underlying dysregulation of erythropoiesis. We previously characterized individuals from both CMS and non-CMS populations and recapitulated the EE phenotype *in vitro* with minimal interclonal and subject variability^5^. Interestingly, we find a remarkable difference in the EE response only under hypoxia between CMS and non-CMS groups as illustrated in our previous studies^5-8^. In this study, to further elucidate the transcriptional and epigenomic drivers of divergent erythropoietic responses in CMS and non-CMS individuals, we performed integrated RNA-seq and ATAC-seq analyses on iPSC-derived erythroid cells differentiated under both hypoxic and normoxic conditions.

### Study Design

Study subjects (3 CMS and 3 non-CMS) were adult males (23-45yrs) born above 4000 m and residing in the Andean mountains (∼4338m). Subjects signed an informed written consent under protocols approved by UC San Diego and Universidad Peruana Cayetano Heredia. CMS patients were defined by high CMS score (17-27) and marked by elevated hematocrit (average: 69%). Erythroid cells were generated under normoxic and hypoxic conditions following our established iPSC-derived Embryoid bodies (EB) *in vitro* platform^5,6,9^. We performed RNA-seq and ATAC-seq on week 3 EBs, which consisted predominantly of reticulocyte-stage erythroid cells (CD71^+^, CD235a^+^), representing a critical point in erythroid differentiation. RNA-seq libraries were generated using Illumina rRNA Depletion and stranded Total RNA kits (PE100). ATAC-seq transposition experiments were performed on 50,000 cells/sample with Illumina TDE1 Enzyme^4^ and libraries generated using the Kapa Biosystems and Illumina kits (PE50). Libraries were sequenced at the UC San Diego IGM Genomics Center (NovaSeq 6000). Sequencing processing was performed as previously described^6,10^: RNA-seq samples were mapped with *kallisto* to the GRCh38.12 v109 transcriptome and differential gene expression (DGE) performed with *limma-voom* (FDR *p*-value <0.05); ATAC-seq samples were aligned to the GRCh38.12 genome with *BBMap* v37.95, ATAC-seq peaks called with *Genrich*, and differential accessibility performed with *csaw* and *edgeR glmQLFit* (FDR *p*-value <0.05). BFU assay was conducted by standard procedure as detailed in supplementary.

## Results and Discussion

We characterized the erythropoietic response of CMS and non-CMS subjects under normoxia and hypoxia by RNA-seq and ATAC-seq (Figure 1A-B). RNA-seq PCA revealed substantial CMS- and hypoxia-associated variance along PC1 (46.7%). K-means clustering revealed clusters enriched for inflammation, hematopoietic and erythroid lineage, and hypoxia response that distinguished CMS subjects under both normoxia and hypoxia (Figure 1D; supplemental Figure 1A-B). Differential Gene Expression analysis identified a substantial number of DEGs in CMS under both normoxia (10229) and hypoxia (6454) (Figure 1E-F). Unique hypoxia DEGs (2024) were enriched for inflammation and vasculature development (sFigure 2A-B). Enrichment analysis revealed activation of hematopoiesis, heme metabolism, and p53 genesets under normoxia and further augmented under hypoxia (Figure 1G). Similarly, WNT/β-catenin and Notch signaling–known repressors of erythroid commitment–were downregulated under normoxia and further suppressed under hypoxia. Importantly, we observed activation of three key inflammatory programs: the stress erythropoiesis-associated EPOR-JAK2-STAT5 (IL2-STAT5 geneset); the IL6-JAK1-STAT3; and the canonical NFκB program (Figure 1H, FDR *p*-value <1×10^−10^). While the JAK-STAT programs were elevated under normoxia and augmented under hypoxia, NFκB activation was hypoxia-specific. PROGENy pathway analysis corroborated this observation, uncovering hypoxia-induced augmentation of JAK-STAT signaling and hypoxia-specific activation of NFκB in CMS (Figure 1I). Additional pathways included sustained VEGF activity and increased androgen signaling, reinforcing previous findings^5,11,12^.

**Figure 1.**
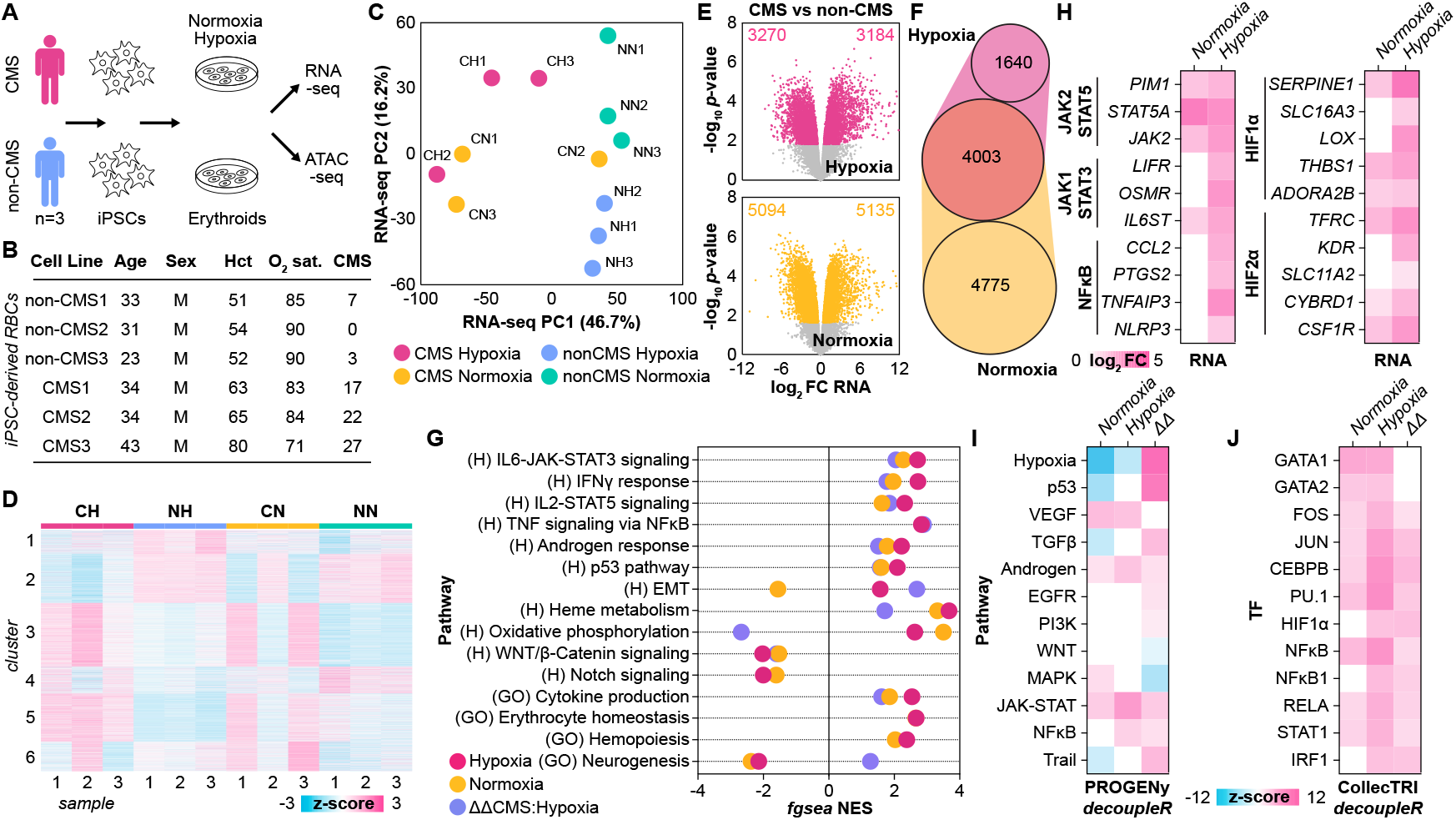
RNA-seq analysis shows a distinct hypoxia-responsive inflammatory signature in CMS erythroid cells. **A** Schematic of the experimental workflow. Patient-derived CMS and non-CMS hiPSCs were differentiated into erythroid cells through generation of EBs and RNA-seq and ATAC-seq was performed at week 3 EB stage which predominantly is comprised of CD71^+^/CD235a^+^ erythroid cells. **B** Demographic and clinical characteristics of study participants, including age, sex, hematocrit, and oxygen saturation. **C** RNA-seq PCA of genes with >100cpm expression. Notably, one subject (CMS2) aligned with non-CMS subjects under normoxia and only exhibited the disease phenotype under hypoxia, supporting the view that CMS forms a polygenic continuum rather than a binary condition. **D** z-score heatmap of clusters identified in RNA-seq data in C. **E** RNA-seq volcano plots of differentially expressed genes (DEGs) CMS relative to non-CMS under hypoxia and normoxia (*limma* FDR *p*-value < 0.05). **F** Venn-Euler diagram of DEGs in E. **G** Gene Ontology: Biological Process (GO) and Hallmark (H) database geneset enrichment using *fgsea*; dot plots indicate significant (FDR *p*-value < 0.05) pathways in CMS vs non-CMS under normoxia, hypoxia, or ΔΔCMS:Hypoxia (ΔHypoxia – ΔNormoxia), revealing activation of inflammatory (IL-6–JAK-STAT3, IFNγ, TNF–NFκB) and stress-erythropoiesis programs in CMS erythroid cells. **H** Canonical targets of JAK2-STAT5, JAK1-STAT3, and NFκB signaling or HIF targets (HIF1α and HIF2α) differentially expressed under hypoxia (FDR *p*-value < 0.05). **I-J** *decoupleR* activity inference using the **I** PROGENy pathway and **J** CollecTRI TF regulon database (FDR *p*-value < 0.05), showing elevated NFκB, STAT, and HIF activity.

**Figure 2.**
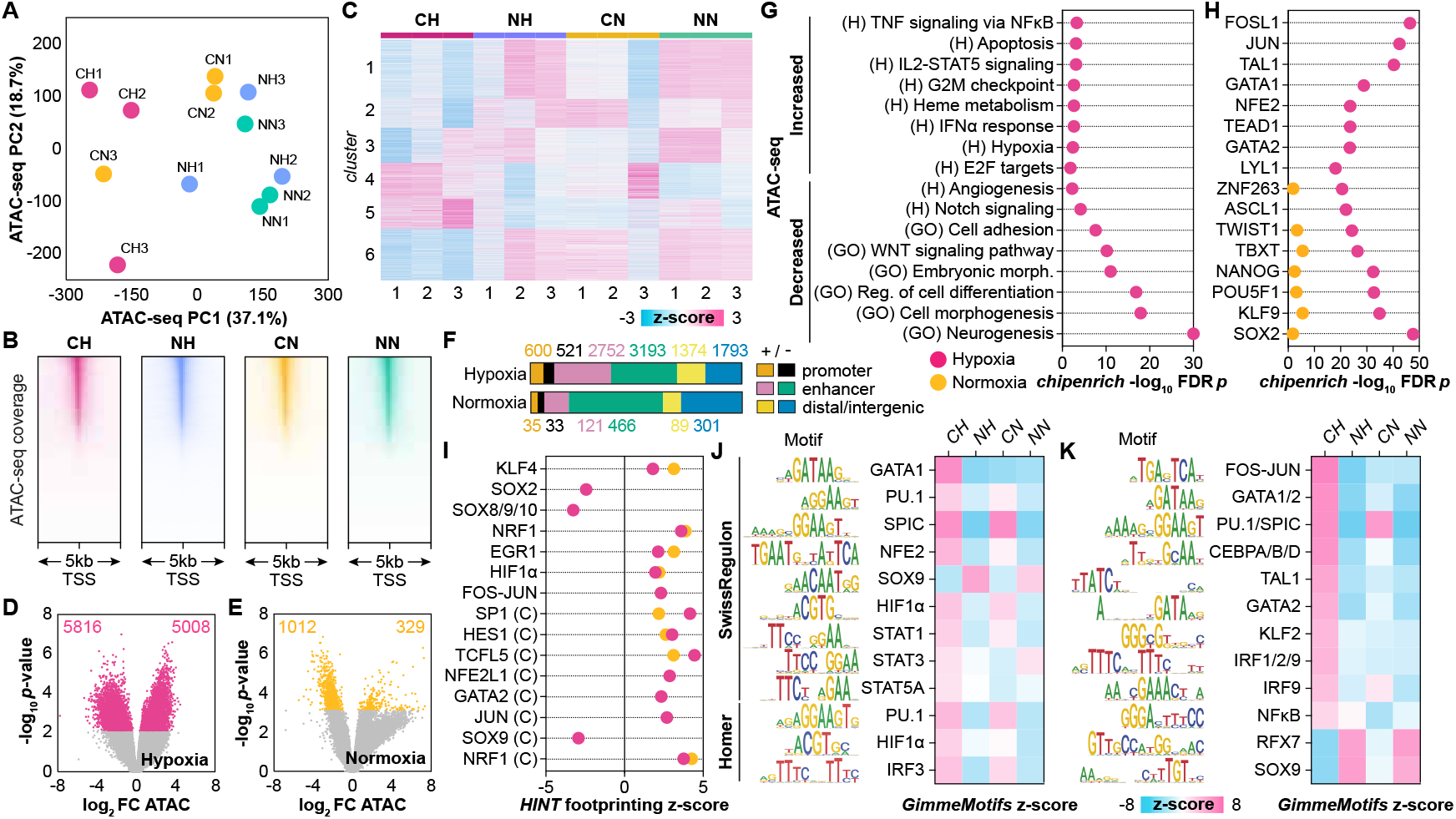
ATAC-seq reveals enhanced chromatin accessibility and inflammatory transcription factor activation in CMS erythroid cells. **A** ATAC-seq PCA of peaks with >100cpm expression. **B** ATAC-seq coverage plots around TSS for all 4 subject types. **C** z-score heatmap of clusters identified in ATAC-seq data in A. **D-E** ATAC-seq volcano plots of differentially accessible regions (DARs) CMS relative to non-CMS under **D** hypoxia and **E** normoxia (*edgeR* FDR *p*-value < 0.05). **G-H** *chipenrich* enrichment analysis of DARs using the **G** Gene Ontology: Biological Process (GO) and Hallmark (H) databases and **H** ReMap TF-gene target database (*chipenrich* FDR *p*-value < 0.05), highlighting strong enrichment of inflammatory and erythropoietic regulators (e.g., NFκB, STAT, FOS-JUN). **I** Top differentially activated and repressed TFs across in CMS under normoxia and hypoxia by HINT TF footprinting analysis using the Swiss Regulon and CISBP (C) databases (*p*-value <0.05). **J-K** TF motif activity in accessible ATAC-seq regions using the **J** Swiss Regulon (SR) and Homer (H) and **K** GimmeMotifs (GM) databases, showing elevated activity of GATA1/2, NFκB, STAT5, and KLF family motifs in CMS under hypoxia.

Aligned with our pathway enrichment results, transcription factor (TF) activity analysis revealed activation of TFs that regulate erythropoiesis and hematopoiesis (GATA1/2, FOS/JUN, TAL1, LYL1, PU.1) and stress response (CEBPB) under normoxia and augmented under hypoxia (Figure 1J; Supplemental Figure 3A). STAT factors involved in JAK-STAT signaling (e.g. STAT1, STAT3, STAT5A) were elevated under normoxia and augmented under hypoxia, while NFκB activation, particularly the NFκB1 subunit of the canonical RelA:p50 heterodimer, was activated more specifically under hypoxia. Interestingly, IRF1, which can be activated by both STAT and NFκB signaling, displayed the same trend. Although *HIF1A* expression was unchanged, predicted HIF1α activity also increased in a hypoxia-specific manner. *EPAS1* (HIF2α) expression was induced by hypoxia, along with key targets of HIF2α, illustrating activation of the hypoxia response through both HIF1α and HIF2α arms. Collectively, these results demonstrate that under normoxia, CMS exists in a heightened inflammatory and erythroid lineage state, primed for hypoxia-selective exacerbation.

**Figure 3.**
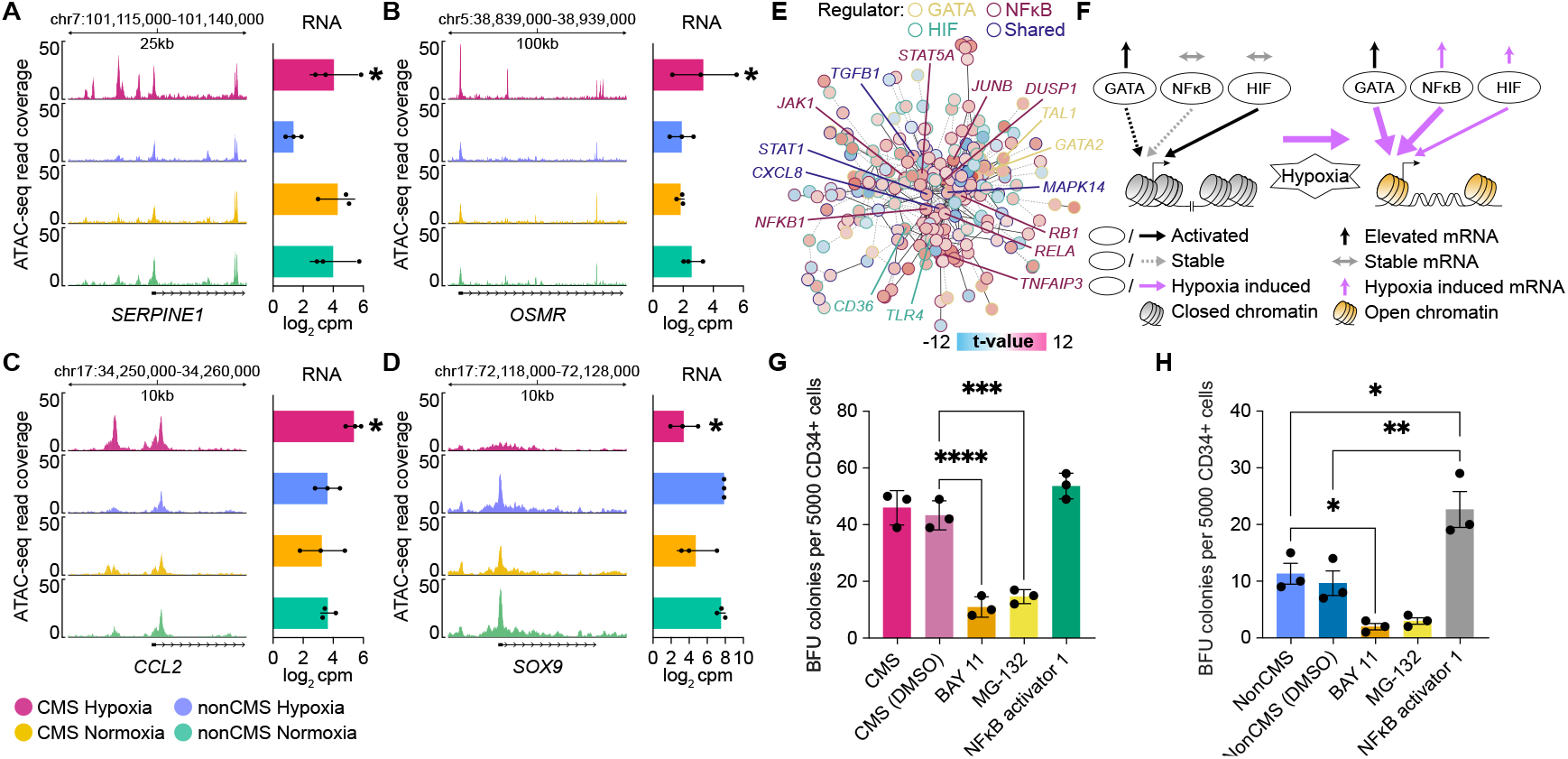
Coordinated activation of GATA, NFκB, and HIF networks drives hypoxia-induced erythroid and inflammatory reprogramming in CMS. **A-D** ATAC-seq coverage plots (left) and RNA-seq expression (right) differential in CMS vs non-CMS under hypoxia for targets of **A** HIF1α/HIF2α (*SERPINE1*), **B** JAK1-STAT3 signaling (*OSMR*), **C** NFκB signaling (*CCL2*), **D** ectoderm lineage (*SOX9*). * = ΔRNA FDR *p*-value <0.05. **E** PPI network of genes regulated by GATA (GATA1/2), HIF (HIF1α, HIF2α), NFκB (RELA, NFKB1), or shared (2 or more TFs), highlighting the shared regulation and StringDB PPI network interaction of hypoxia-responsive gene programs in CMS. Black edge, intra-network PPI interaction; grey dotted edge, inter-network PPI interaction. Colored by *limma* differential *t*-value in CMS under hypoxia. **F** Key regulatory mechanisms in CMS under normoxia and hypoxia. While JAK1 and JAK2 signaling is upregulated constitutively in CMS under normoxia and augmented under hypoxia, GATA, NFkB, and HIF factor function is induced in a hypoxic-specific manner. GATA factor expression is elevated in normoxia as are GATA targets, but GATA TF motif activity and target gene chromatin accessibility is induced by hypoxia. NFκB, however, is activated— RNA expression, predicted TF activity, and TF motif activity—specifically under hypoxia. Finally, while HIF footprinting suggests TF motif binding under normoxia, hypoxia activates HIF mRNA expression and TF motif activity, leading to HIF target gene expression. **G-H** BFU-E colony formation in **G** CMS or **H** non-CMS cells treated with no inhibitor, DMSO, NFκB inhibitor (BAY 11 and MG-132) or NFκB Activator 1 (HY-134476). *n*=3 for each experiment; * denotes *p*-value <0.05, ** *p*-value <0.01, *** *p*-value <0.001 and **** *p*-value <0.0001.

ATAC-seq analysis revealed comparable genomic coverage across all subjects and PCA positioned hypoxia response on PC1 (37.1% variance) (Figure 2A-B). K-means clustering identified clusters enriched for inflammation, hypoxia response, and erythroid lineage state distinguishing CMS subjects, particularly under hypoxia (Figure 2C; Supplemental Figure 4A-B). Differentially accessible region (DAR) analysis revealed a striking increase in hypoxia-induced chromatin changes in CMS, with 10,824 DARs identified under hypoxia versus 1,341 under normoxia (Figure 2D), primarily at enhancer regions (Figure 2F). Enrichment analysis of DARs revealed increased accessibility at genes related to inflammation, heme metabolism, hypoxia, and apoptosis (Figure 2G) as well as targets of erythroid lineage regulators (e.g., FOS/JUN, GATA1/2, TAL1) under hypoxia (Figure 2H). Concordant with the RNA-seq, hypoxia repressed genes associated with ectodermal lineage (GOBP Neurogenesis, ASCL1 targets), pluripotency (SOX2, NANOG), and WNT/B-Catenin and Notch signaling in CMS. TF footprinting (*HINT-ATAC*) and activity (*GimmeMotifs*) analysis revealed hypoxia-specific chromatin accessibility changes critical to CMS pathogenesis (Figure 2I-K): GATA and FOS-JUN TFs were unchanged in normoxia yet robustly activated in hypoxia, indicating that their TF activity is driven by both occupancy and chromatin accessibility change rather than merely TF expression. In contrast, HIF1α footprinting was similarly elevated under normoxia and hypoxia, while motif activity was augmented under hypoxia (Figure 2G,J). This suggests a chromatin priming model, where pre-established HIF1α occupancy enables hypoxia to enhance accessibility at poised sites, driving hypoxia-specific target gene expression in CMS. TF motif activity of JAK-STAT signaling factors was elevated under normoxia (e.g., STAT1, STAT3, STAT5A) and further augmented under hypoxia (Figure 2J), as observed in RNA-seq. Conversely, IRF1 and NFκB showed minimal activity under normoxia but were strongly activated under hypoxia (NFκB *GimmeMotifs* ΔΔz-score 4.51), aligning with the NFκB-driven inflammation response observed in RNA-seq (Figure 2K). While ETS-family TFs (e.g., PU.1, SPIC) remained consistently elevated, the erythroid-specific FOS-JUN and GATA family factors (GATA1/2, TAL1) were selectively activated under hypoxia by both TF footprinting and motif activity analysis (ΔΔz-score 12.43 and 8.19, respectively). Finally, we observed the decreased and hypoxia-inducible repression of TF activity and footprinting of pluripotency (SOX2, ΔΔz-score −1.82) and ectoderm lineage (SOX9, ΔΔz-score −5.13) factors, likely driven by decreased occupancy at motif sites as evidenced by loss of TF footprinting. Key markers for HIF response (*SERPINE1*), JAK-STAT signaling (the JAK1-STAT3 upstream receptor *OSMR*), NFKB signaling (*CCL2*), and ectoderm lineage state (SOX9) were differentially activated (*SERPINE1, OSMR, CCL2*) or repressed (*SOX9*) in CMS specifically under hypoxia (Figure 3A-D). Finally, we integrated TF-gene regulation and StringDB PPI interaction for the core hypoxia-induced GATA, NFκB, and HIF programs (Figure 3E). This revealed essential genes involved in erythropoiesis as NFκB-controlled at the network center, including *MAPK14, DUSP1*, and *CXCL8*, highlighting functional overlap between these three programs. Overall, our integrative multiomics reveals that in CMS, GATA, and HIF programs are primed under normoxia and, together with NFκB, are aberrantly activated upon hypoxia exposure (Figure 3F). While HIF^13,14^ and GATA1^5,6^ have been linked to erythropoiesis in Andean highlanders, NFκB-mediated inflammation has not been studied in this context.

To establish a causal role for hypoxia-specific, NFκB-mediated inflammation in driving excessive erythropoiesis, we functionally modulated NFκB signaling in erythroid cells and assessed BFU-E colony formation (Figure 3G-H). Inhibition of NFκB led to a striking reduction in BFU-E colonies specifically in CMS (*p*-value <0.0001), indicating their dependency on inflammatory signaling. Conversely, activation of NFκB in non-CMS cells nearly doubled BFU-E colony numbers (*p*-value <0.01), emphasizing the potential of this pathway to enhance erythropoiesis under hypoxic conditions.

Our study presents the first multiomics framework that elucidates the disease-specific mechanisms underlying excessive erythropoiesis (EE) in CMS (Monge’s disease). We demonstrate that CMS erythroid cells exist in a primed state for EE even under normoxia, indicating an intrinsic predisposition. Hypoxia amplifies this state, demonstrating how nvironmental exposure can unmask latent vulnerabilities (Figure 3F). This mirrors stress erythropoiesis, but in CMS the response is chronically engaged, driving pathological EE. Importantly, NFκB-mediated inflammatory signaling emerges as an active driver rather than a secondary effect, positioning inflammation as a central therapeutic target in high-altitude–associated erythropoietic disorders.

## Data Sharing Statement

RNA-seq and ATAC-seq data are available at the NCBI GEO under the accessions GSE309468 and GSE309467, respectively. Normalized count matrices for RNA-seq and ATAC-seq data are provided in Supplementary files.

## Acknowledgements

We sincerely thank Ms. Orit Poulsen and Daniela Bermudez for their assistance and the IGM Core at UCSD for sequencing and technical assistance.

## Authorship and conflict-of-interest statements

Conceptualization: P.A., G.G.H., D.Z.

Subject recruitment: F.C.V.

iPSC line and RBC generation: P.A., S.B.

Sequencing Experiments: P.A.

Data Analysis: A.B.C.

Visualization: A.B.C.

Supervision: G.G.H., S.S.

Writing—original draft: P.A., A.B.C.

Writing—review & editing: P.A., A.B.C., F.C.V., S.B., D.Z., S.S., G.G.H.

Conflict-of-interest disclosure: The authors declare no competing financial interests.

