## supplementary information for "Integrated multiomics reveals inflammation-driven excessive erythrocytosis in subjects with Monge’s disease"

Azad *et al.* 2025

*Supplementary Methods*

### iPSC Characterization and Erythroid Differentiation

Human iPSCs were generated from fibroblasts, confirmed for pluripotency by DNA fingerprinting, karyotype analysis, pluripotency marker expression, and trilineage differentiation, and subsequently expanded on Matrigel. Erythroid differentiation was induced using a modified Kobari et al. (2012) protocol, which we have optimized and validated in our previous studies based on stage-specific CD markers, hemoglobinization, and overall differentiation efficiency under normoxia and hypoxia. In brief, human iPSCs were expanded on Matrigel-coated dishes (BD Biosciences) in mTeSR medium (STEMCELL Technologies). Differentiation toward the erythroid lineage was carried out following a modified protocol based on Kobari et al. (2012). For each subject, cultures were initiated with approximately 1×10^7 cells. Embryoid bodies (hEBs) were formed and maintained for 27 days in liquid IMDM (Biochrom) supplemented with 450 µg/ml holo-human transferrin (Sigma-Aldrich), 10 µg/ml recombinant human insulin (Roche), 2 IU/ml heparin, and 5% human plasma. The culture medium additionally contained 100 ng/ml SCF, 100 ng/ml TPO, 100 ng/ml FLT3 ligand, 10 ng/ml BMP4, 5 ng/ml VEGF-A165, 5 ng/ml IL-3, 5 ng/ml IL-6 (PeproTech), and 3 U/ml erythropoietin.

### RNA-seq data processing and analysis

RNA was isolated from CMS and non-CMS ($n = 3$ subjects/condition) hiPSC-derived erythroid cells grown under hypoxic and normoxic conditions for two weeks using the Zymo R1050 RNA Kit. RNA-seq libraries were prepared using the Illumina Ribo-Zero Plus rRNA Depletion and TruSeq Stranded Total RNA Library Prep Kits (paired-end 100bp reads). RNA-seq libraries were sequenced on a NovaSeq 6000 generating paired-end, 100 bp (PE100) reads at the UC San Diego IGM Genomics Center. Preprocessing of RNA-seq data was performed using the TrimGalore! Package v0.6.4, removing adapter sequences and low-quality reads using CutAdapt v1.18. Trimmed RNA-Seq reads were mapped to the GRCh38.12 v109 human transcriptome using kallisto v0.50 followed by transcript level summation to the gene level using the R package *tximport* v1.36. Lowly expressed genes were filtered out using *filterByExpr* and counts normalized using the weighted mean trimmed of M-values (TMM) in the R package *edgeR* v4.6.3. Differential gene expression (DGE) analysis was performed using the *voomwithQualityWeights* function within the *limma* v3.64 R package, defining differentially expressed genes (DEGs) with a false discovery rate (FDR)-adjusted *p*-value cutoff of $< 0.05$ using the Benjamini–Hochberg correction method from the *eBayes* differential t-test. Quasi-proportional Venn diagrams of DEG overlap between the CMS vs non-CMS under hypoxia and normoxia were generated using the *nVennR* v0.2.3 R package. Clusters in RNA-seq data were identified in genes passing a threshold of > 100cpm across all samples using the *kmeans* function in the R *stats* package. Geneset enrichment of clusters was carried out using hypergeometric enrichment (*tmodhgtest*) in the *tmod* R package with the Hallmark and Gene Ontology Biological Process (GOBP) geneset databases. Geneset enrichment analysis was performed using the *fgsea* R package with the Hallmark and GOBP geneset databases and the ENCODE-ChEA Consensus and ReMap TF-gene target databases. Pathway level activity analysis was performed using the *decoupler mlm* algorithm with the PROGENy pathway database, while transcriptional factor (TF) activity analysis was performed using *decoupler ulm* algorithm with the CollecTRI TF regulon database. For all *fgsea* and *decoupler* tests, genes were ranked by the *limma* t-value.

### ATAC-seq data processing and analysis

ATAC-Seq transposition experiments were performed on 50,000 CMS and non-CMS ($n = 3$ subjects/condition) hiPSC-derived erythroid cells grown under hypoxic and normoxic conditions for two weeks. Cells were washed with PBS, lysed with cold lysis buffer (10 mM Tris-HCl, pH 7.4, 10 mM NaCl, 3 mM MgCl2, 0.1% IGEPAL CA-630), and suspended in 50 µl of 1X reaction buffer (25 µl of Tagment DNA Buffer, 2.5 µl of Tagment DNA enzyme I, and 22.5 µl of water) (Illumina, Cat. 15028523) as previously described (Buenrostro et al. 2013). Transposase reactions were carried out at 37 °C for 30 min, and DNA was purified using ChIP DNA Clean & Concentrator kits (Zymo Research, Cat. D5205). Libraries were generated using the Kapa Biosystems Real-Time Library Amplification Kit. The resulting libraries were sequenced on an Illumina Novaseq 6000 platform generating paired-end, 50 bp (PE50) reads with an average of over 100 million reads per sample. ATAC-Seq data preprocessing was performed with TrimGalore!, removing sequencing adaptors and selecting all paired-end reads above a quality score threshold (Phred $Q > 20$). Trimmed reads were aligned to the GRCh38.12 genome with BBMap v37.95 in the BBtools suite with the options maxindel = 20 ambig = random, followed by sorting and indexing of bam files using Samtools v1.35 and annotation of PCR duplicates using the Picard v2.3.0 markDuplicates function. Genrich v0.61 was used to call peaks on the ATAC-Seq data to determine regions of accessible chromatin. Consensus peaks (n=2 per condition) were generated using *Diffbind* v3.18.0 and peak coverage calculated with *csaw.* Loess normalization was performed across peak windows and differential peaks for each comparison of interest determined using *edgeR* glmQLFit with an FDR *p*-value cutoff of 0.05. Differentially Accessible Regions (DARs) were annotated using *ChIPseeker*, defining the promoter region -2000 to +500 bp from the TSS. Enhancer-associated ATAC-Seq regions were defined as differential peaks occurring within the region list of TSS-associated enhancers generated by the FANTOM5 project using the join_overlap_inner function in the *plyranges* R package. ATAC-seq coverage heatmaps were generated with the deepTools functions *computeMatrix* and *plotHeatmap*. Clusters in ATAC-seq data were identified in peaks with high variance (var > 0.5) across all samples using the *kmeans* function in the R *stats* package. Geneset enrichment of clusters was carried out using hypergeometric enrichment (*tmodhgtest*) in the *tmod* R package with the Hallmark and Gene Ontology Biological Process (GOBP) geneset databases. To identify candidate ontologies and transcription factors (TFs) potentially associated with CMS, we performed enrichment analysis of DARs using the logistic regression model test function chipenerch in the *chipenrich* R package with the GO:BP and Hallmark (ontology) or ReMap (TF) databases. HINT-ATAC differential footprinting motif activity analysis between the CMS and non-CMS under normoxia and hypoxia was performed with the SwissRegulon and CIS-BP motif databases (*p*-value < 0.05) using the Regulatory Genomics Toolbox. GimmeMotifs *maelstrom* differential motif activity was performed for the same comparisons using the SwissRegulon, GimmeMotifs (based on CIS-BP), and Homer motif databases.

**Network analysis**

To generate TF-gene and PPI integrative networks of GATA, NFkB, and HIF programs in CMS under hypoxia exposure, we curated targets of GATA (GATA1 and GATA2 targets), NFκB (RELA and NFκB1 targets), and HIF (HIF1α and HIF2α) from the CollecTRI TF-gene regulon database and subset for DEGs. We subset each group of DEGs into GATA, NFκB, HIF, or shared (NFκB + GATA or HIF control) and projected PPI interactions within each group (solid black line) or across each group (dotted grey line) using the StringDB PPI database.

**Functional assay (colony assay) with inhibitors**

Experiments were performed with the inhibitors using the following concentrations in the colony forming assays using iPSC-derived CD34+ cells as described below: BAY 11-7082(5µM), MG-132( 5µM), NF-κB activator 1 (1 µM) and MCC950 (10ug). After 7 days of iPSC differentiation, EBs were dissociated with Accutase and $CD34^+$ cells purified using EasySep selection (STEMCELL Technologies). Purified $CD34^+$ cells were plated ($1\times10^5$/dish) in MethoCult (H4034) with 2% FBS under hypoxia (5% $O_2$, 14 days) for BFU-E, CFU-E, and CFU-GEMM scoring.

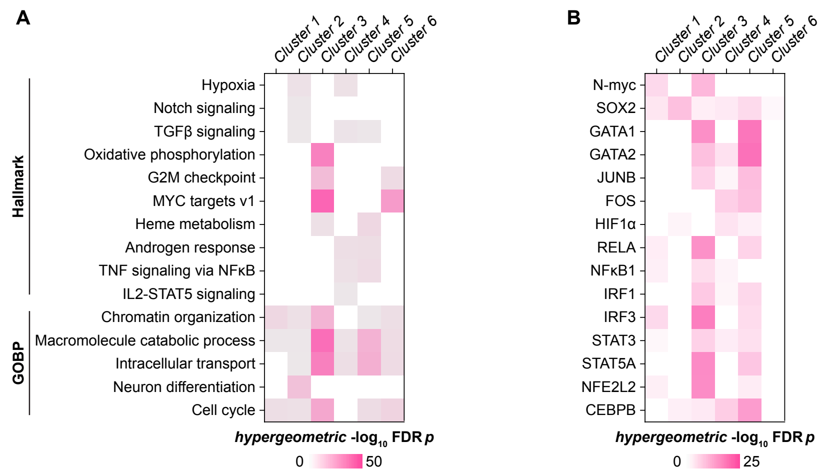


**Supplemental Figure 1**

**A-B** *tmod* hypergeometric enrichment of genes in the 6 clusters identified by k-means clustering of RNA-seq data using the **A** Gene Ontology: Biological Process and Hallmark databases geneset enrichment or **B** ReMap TF-gene target database.

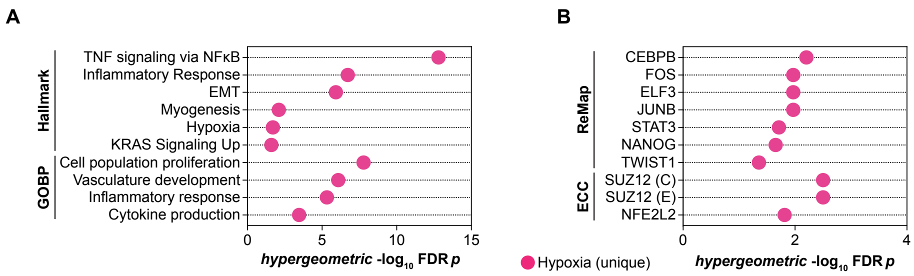


**Supplemental Figure 2**

**A-B** *tmod* hypergeometric enrichment of DEGs unique to CMS under hypoxia using the **A** Gene Ontology: Biological Process and Hallmark databases geneset enrichment or **B** ENCODE-ChEA Consensus (ECC) or ReMap TF-gene target databases.

A

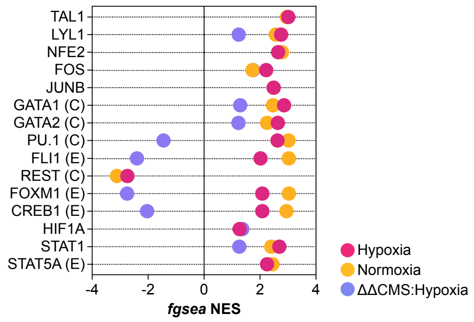


**Supplemental Figure 3**

**A** *fgsea* TF geneset enrichment using the ENCODE-ChEA Consensus (ECC; ENCODE, (E); ChEA, (C)) or ReMap TF-gene target databases; dot plots indicate significant (FDR *p*-value < 0.05) pathways in CMS vs non-CMS under normoxia, hypoxia, or ΔΔCMS:Hypoxia (ΔHypoxia – ΔNormoxia).

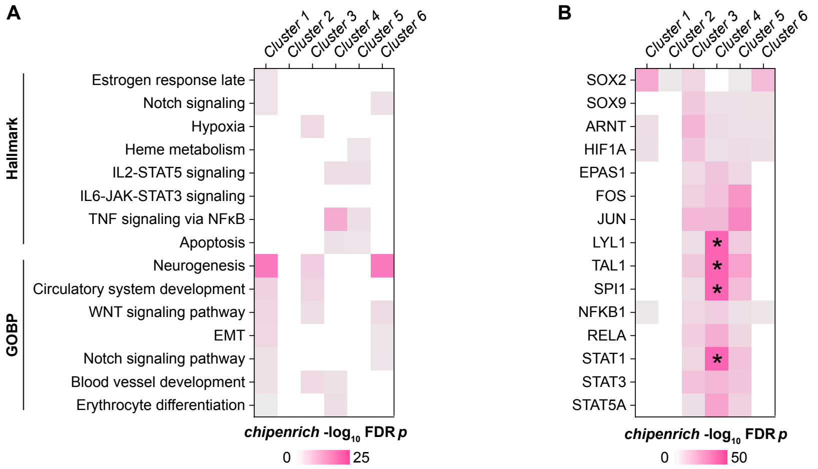


### Supplemental Figure 4

**A-B** *chipenrich* enrichment analysis of genes in the 6 clusters identified by k-means clustering of ATAC-seq data using the **A** Gene Ontology: Biological Process (GO) and Hallmark (H) databases and **B** ReMap TF-gene target database (*chipenrich* FDR $p$-value $< 0.05$; *=FDR $p$-value $< 1 \times 10^{-50}$).
